# Production and analysis of a mammalian septin hetero-octamer complex

**DOI:** 10.1101/2020.06.15.152751

**Authors:** Barry T. DeRose, Robert S. Kelley, Roshni Ravi, Bashkim Kokona, Elias T. Spiliotis, Shae B. Padrick

**Affiliations:** Department of Biochemistry and Molecular Biology, Drexel University, Philadelphia, PA 19102; VCU Health System, Richmond, VA, 23219; WuXi Advanced Therapies, Philadelphia, PA 19112; Department of Chemistry, Haverford College, Haverford PA, 19041-1391, USA; Department of Biology, Drexel University, Philadelphia, PA 19104

**Keywords:** Protein complex, cytoskeleton, septins, polymerization, biochemical reconstitution, GTP-binding proteins

## Abstract

The septins are filament-forming proteins found in diverse eukaryotes from fungi to vertebrates, with roles in cytokinesis, shaping of membranes and modifying cytoskeletal organization. These GTPases assemble into rod-shaped soluble hetero-hexamers and hetero-octamers in mammals, which polymerize into filaments and higher order structures. While the cell biology and pathobiology of septins are advancing rapidly, mechanistic study of the mammalian septins is limited by a lack of recombinant hetero-octamer materials. We describe here the production and characterization of a recombinant mammalian septin hetero-octamer of defined stoichiometry, the SEPT2/SEPT6/SEPT7/SEPT3 complex. Using a fluorescent protein fusion to the complex, we observed filaments assembled from this complex. In addition, we used this novel tool to resolve recent questions regarding the organization of the soluble septin complex. Biochemical characterization of a SEPT3 truncation that disrupts SEPT3-SEPT3 interactions is consistent with SEPT3 occupying a central position in the complex while the SEPT2 subunits are at the ends of the rod-shaped octameric complexes. Consistent with SEPT2 being on the complex ends, we find that our purified SEPT2/SEPT6/SEPT7/SEPT3 hetero-octamer copolymerizes into mixed filaments with separately purified SEPT2/SEPT6/SEPT7 hetero-hexamer. We expect this new recombinant production approach to lay essential groundwork for future studies into mammalian septin mechanism and function.

## Introduction

Septins are a family of GTP-binding proteins that are found in eukaryotic cells. They were first discovered in the budding yeast *Saccharomyces cerevisiae* during studies of cell division cycle (CDC) gene mutants.^1^ A family of CDC proteins essential for cytokinesis were found localized as rings at the bud neck and were named septins.^2^ Orthologous proteins have since been found in mammalian cells.^3^ In addition to a key role in cytokinesis,^4–6^ mammalian septins have been implicated in important cellular processes, including membrane trafficking,^4,7–12^ acting as diffusion barriers,^13–15^ stress fiber assembly,^16–20^ cellular adhesions^16,21^ and intracellular pathogen invasion and spread.^22–27^ Beyond their roles in normal cell biology, their dysfunction has been implicated in heritable diseases^28,29^ and numerous cancers.^30–33^ Despite this growing importance, the biochemical mechanisms underlying these roles are not well understood.

All septins have a core GTP-binding domain with sequence similarity of at least 70%, an N-terminal helix enriched in basic amino acids and a C-terminal highly conserved sequence of 53 amino acids called the septin unique element (SUE).^3,7^ In humans, septin proteins are encoded by thirteen different genes: SEPT1 through SEPT12 and SEPT14.^34^ These are classified into four sub-groups which are defined by sequence homology: SEPT2, SEPT3, SEPT6, and SEPT7.^7,35,36^ These diverse septin subunits organize into soluble complexes that are predominantly hetero-hexamers or hetero-octamers, with assembly being guided by two stereotypical interactions. Two surfaces on opposite sides of the septin monomer interact with other septin monomers. The first surface overlaps the GTP-binding site (G-interface) and the second incorporates elements at their amino and carboxy termini (N-/C-interface).^37–40^ The complex assembly is then controlled by interface dependent binding preferences, e.g. SEPT6 subgroup members prefer to bind SEPT7 via their N-/C-interfaces.^41–44^ Different members of a subgroup can replace each other in a hetero-octamer or hetero-hexamer.^34^ Thus, there are two prototypical complexes, SEPT2/SEPT6/SEPT7 hetero-hexamers and SEPT2/SEPT6/SEPT7/SEPT3 hetero-octamers, and many possible complexes that follow these prototypical organizations.^35,41,45^ While the pairwise interactions are fairly well understood, the overall organization of the soluble complexes is an active and open question. Historically, SEPT2 has been placed at the center of models of these rod-like complexes,^37,41,46^ but recent work has indicated that the SEPT2 subgroup members may in fact be on the ends.^45,47^

These soluble hexamers and octamers then assemble into higher order structures such as filaments, bundles, and rings.^17,48^ Whether the filaments in these higher order structures are composed of a heterogenous mix of hetero-hexamers and hetero-octamers, or if distinct hexamer and octamer filament systems exist, has not yet been determined. Notably, the two proposed soluble complex models make opposite predictions, with the old model predicting that hexamers and octamers cannot co-assemble, while the new model predicts they will coassemble. Understanding of these mechanistic differences is important as the assembly and association of these higher order components with cell membranes, actin filaments, and microtubules are fundamentally tied to septin cellular function.^24,49–55^

Septin hetero-hexamers and hetero-octamers both form in mammalian cells with the heterooctamers typically being the dominant species.^41,42^ While the cell biology of mammalian septins is advancing rapidly (see above) and the biophysical basis of yeast septin assembly is progressing,^56–61^ biochemical and biophysical studies of mammalian septins have lagged due to the lack of a mammalian hetero-octamer purification. Although some protocols are established for purifying yeast hetero-octamers,^56,57,62^ mammalian hetero-hexamers^17,37,45,63^ and mixed endogenous mammalian octamers,^47,52^ there is currently no strategy for preparation of recombinant mammalian hetero-octamers of well-defined composition. Recombinant methods will be needed to dissect the many roles of individual septins in the context of the heterooctamer and to enable mechanistic study of septin biochemical function through mutagenesis.

Here, through heterologous expression, we produce a polymerization competent SEPT2/SEPT6/SEPT7/SEPT3 complex, which we refer to as SEPT2/6/7/3. This prototypical complex extends previous co-expression strategies for mammalian hetero-hexamers by including SEPT3. SEPT3 was chosen as a first representative SEPT3 subgroup member both for its role in neuronal biology^64^ and lack of a potentially complicating N-terminal extension (as seen on SEPT9, the more widely expressed SEPT3 subgroup member).^44^ It is hoped that future work can build upon this prototypical complex to produce the variety of mammalian septin complexes observed in across cell types. Using this strategy, we present new evidence that SEPT3 is found in the middle of the octamer, supporting recent findings with other septin complexes.^45,47^ Finally, we demonstrate that our SEPT2/6/7/3 hetero-octamers co-polymerize into filaments with mammalian hetero-hexamers, indicating that mixed hexamer-octamer filaments are possible.

## Results and Discussion

### Production and validation of mammalian septin octamers

Yeast septin hetero-octamers^56,57,62^ and mammalian septin hetero-hexamers^37,45,63^ have been produced through co-expression in bacteria. In addition, mammalian SEPT3 family members express in bacteria.^28,40,65^ Thus, we reasoned that bacterial co-expression of all four subunits may enable complex formation. We focused on a SEPT3 containing complex for this study. We assembled coding sequences for SEPT3 and SEPT2 into a co-expression plasmid (Supplemental Figure 1A) with an origin and antibiotic resistance compatible with an existing SEPT6/SEPT7 co-expression plasmid.^63^ In addition, there is a TEV protease cleavable aminoterminal MBP fusion to SEPT3 and His6 fusion to SEPT2. Thus, amylose affinity chromatography followed by immobilized metal affinity chromatography will obtain septin complexes containing both SEPT3 and SEPT2 (Figure 1A). As field dogma holds that SEPT2 and SEPT3 family members interact via contacts to SEPT6 and SEPT7,^41,66^ respectively, complexes purified by this strategy should contain all four subunits. Following expression and purification, bands consistent with all four septin subunits (including tags) are observed through the double affinity purification (Supplemental Figure 1B).

**Figure 1:**
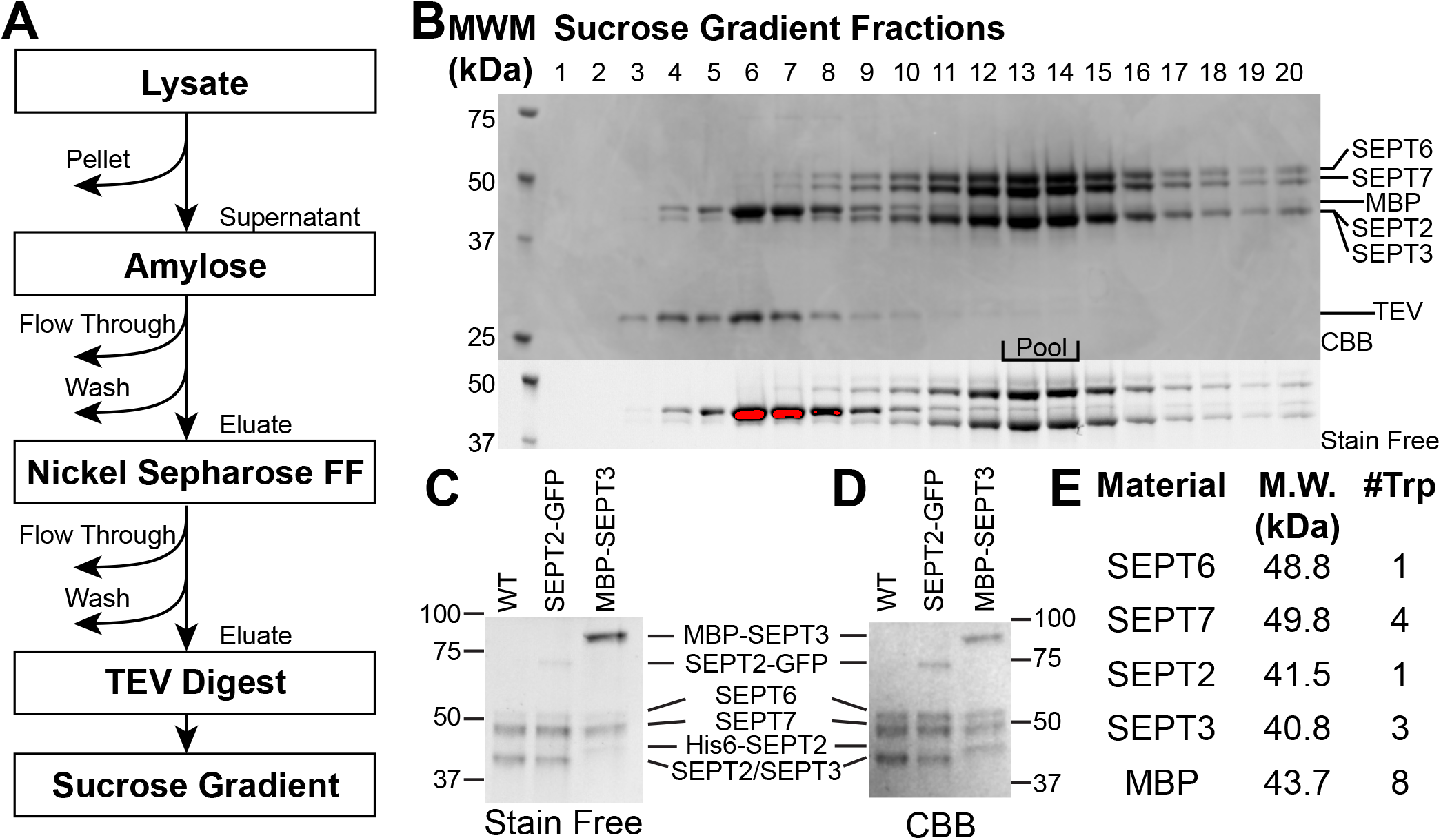
Preparation of SEPT2/6/7/3 complexes. **(A)** Workflow for purification of SEPT2/6/7/3 complexes. Retained fractions are indicated by downward arrows. **(B)** SDS-PAGE analysis of sucrose gradient purification. Fractions (from top of gradient) are indicated across the top of the gel, relevant band mobilities shown at right and molecular weight markers at left. Top subpanel is the Coomassie brilliant blue (CBB) stained gel showing co-migration of four separate bands. Bottom subpanel is the same gel analyzed using ‘Stain Free’ method. Pooled fractions are indicated. Red pixels are overloaded pixels from tryptophan rich MBP in Stain Free imaging. **(C, D)** Identification of bands by relative mobility and staining. MBP-SEPT3/His6-SEPT2 complex is the wild type complex purified without TEV protease treatment. **(E)** Table of predicted molecular weights and tryptophan counts for relevant proteins.

To establish the presence of all four subunits in a complex, we sought a means of separating stable complexes from weakly associated materials. Following proteolytic release of the affinity tags, we used sedimentation into sucrose gradients to resolve the complex from MBP and TEV protease (Figure 1B). There was clear evidence of a multi-subunit complex that sedimented considerably faster than expected for septin monomers (which should sediment similarly to MBP). To identify the subunits co-sedimenting in this complex we used SDS-PAGE to separate and identify the complex composition (Figure 1C-E). To identify these bands, we used the predicted subunit molecular weight and compared band intensity in the ‘Stain Free’ method (Figure 1C) to band intensity with Coomassie Brilliant Blue (CBB) visualization (Figure 1D). The ‘Stain Free’ method is sensitive to tryptophan content,^67,68^ while CBB is more sensitive to total protein content (Figure 1C and D). SEPT6 and SEPT7 migrate more slowly on SDS-PAGE, due to their larger molecular weight. They can be distinguished by their different tryptophan content; SEPT6 has one tryptophan while SEPT7 has four (Figure 1E). The top band is less intense with respect to the ‘Stain Free’ imaging and thus has fewer tryptophan residues. We conclude this slower migrating band is SEPT6, while the other more ‘Stain Free’ intense band is SEPT7. SEPT2 and SEPT3 have very similar molecular weights and a single band appears at the predicted molecular weight for the two. To confirm the presence of both SEPT2 and SEPT3, we tagged SEPT2 with a GFP derivative or left the MBP-SEPT3 and His6-SEPT2 fusions intact. For each of these complexes, similar co-sedimentation was observed (Supplemental Figure 2A and B). These tags increased subunit weight, allowing identification of SEPT2-GFP/SEPT6/SEPT7/SEPT3 and His6-SEPT2/SEPT6/SEPT7/MBP-SEPT3 complexes using mobility on SDS-PAGE (Figure 1C/D/E). Moreover, the CBB staining reveals similar band intensity for each subunit of these tagged complexes, indicating a 1:1:1:1 stoichiometry. In contrast, for the wild type complex the CBB stained SEPT2/SEPT3 band is more intense than the SEPT6 or SEPT7 bands, consistent with additional mass present from two subunits. Thus, we have produced a complex containing members of all four mammalian septin subgroups.

To distinguish a SEPT2/6/7/3 tetramer from higher order complexes, we quantified the sedimentation in our sucrose gradients. Sedimentation rate is sensitive to the solution density and viscosity, as well as particle mass and shape. As particles sediment through the sucrose gradient, they will experience varying solvent density and viscosity as the sucrose concentration increases. As a consequence, directly relating sucrose gradient fraction to sedimentation coefficient under standard conditions (20°C in water) is challenging. However, most proteins have nearly the same partial specific volume, meaning that the changing solution conditions will affect most proteins equally (transmembrane and heavily glycosylated proteins being notable exceptions). Thus, we can measure sedimentation of protein standards in a matched gradient to calibrate the sucrose gradients. We calibrated sedimentation in our gradients using a novel series of blue fluorescent protein standards (Supplemental Figure 3A). These standards were separately analyzed by sedimentation velocity analytical ultracentrifugation (SV-AUC) to measure their sedimentation coefficients under standard conditions of 20°C in water (Supplemental Figure 3B-C). This allowed us to correlate sedimentation fraction to sedimentation coefficient (Supplemental Figure 3D-E). These standards were developed for this experiment and to allow facile detection of the calibration standards in sucrose gradient fractions by fluorescence intensity. These may be broadly useful for sucrose gradient analysis, particularly when analyzing fluorescent protein fusions to targets of interest in complex mixtures.

We applied this calibration to analyze the sedimentation of the SEPT2/6/7/3 complex in our purification gradients. Sedimentation of the SEPT2/6/7/3 wild type complex was measured by integrating the combined SEPT6 and SEPT7 bands from ‘Stain Free’ SDS-PAGE analysis of the sucrose gradient fraction and finding a best estimate of the peak of the fraction profile (Supplemental Figure 3F). For three independent octamer preparations, we analyzed the sucrose gradient purification step to obtain SEPT2/6/7/3 complex sedimentation coefficients of 10.1 S, 10.5 S and 11.3 S (Figure 2A) with an average of 10.6 S.

**Figure 2:**
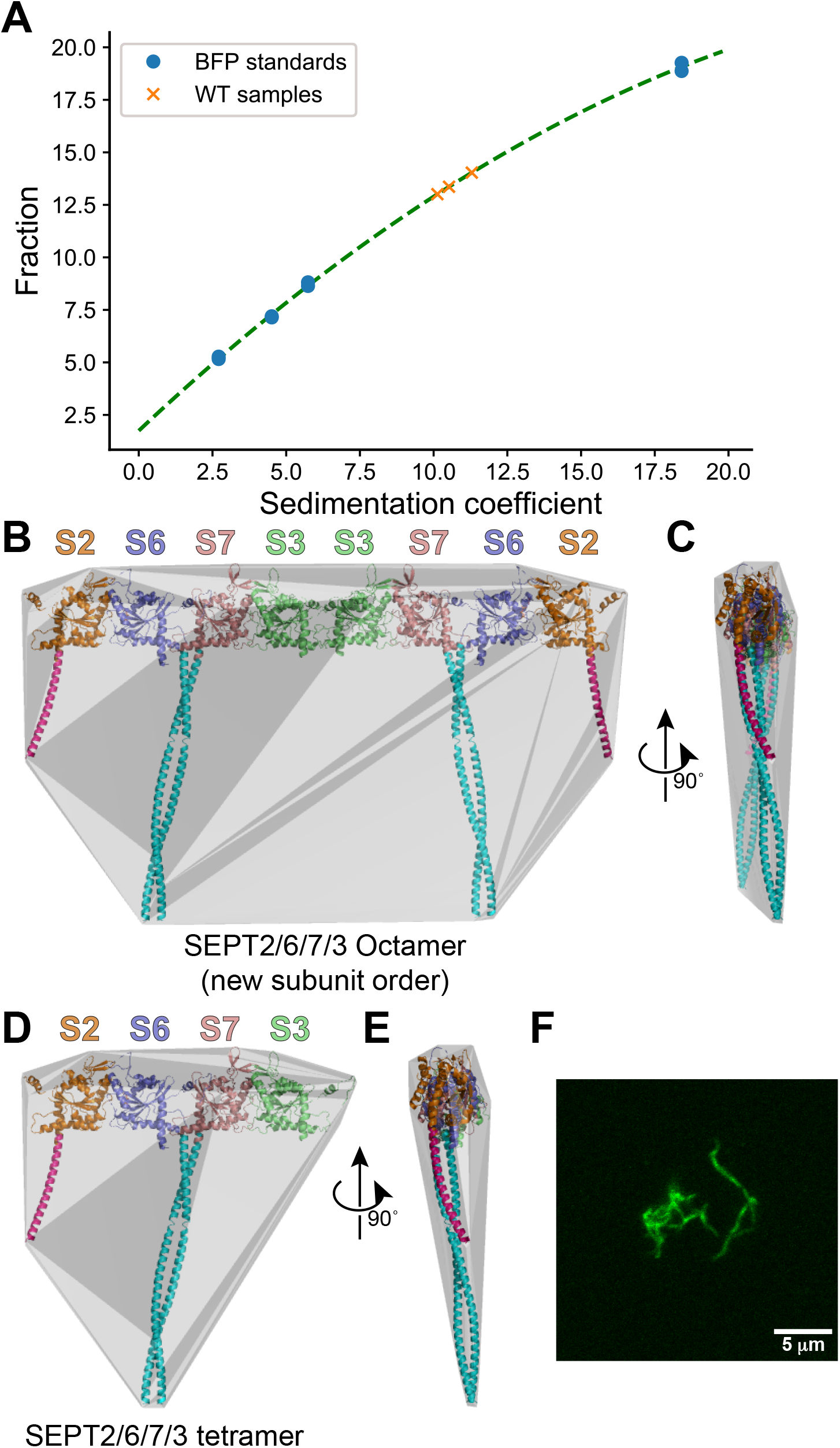
Assessment of SEPT2/6/7/3 complex. **(A)** Sedimentation of SEPT2/6/7/3 complexes into 10-30% w/v sucrose gradients. Sedimentation of complexes is compared to the sedimentation of BFP tagged standards of known sedimentation coefficients (at 20°C in water), with standard curve fit (green dashed line, curve model is a parabola). **(B, C)** SEPT2/6/7/3 octamer model with convex hull (gray surface), with rotation. **(D, E)** SEPT2/6/7/3 tetramer model with convex hull (gray surface), with rotation. **(B-E)** S2, S6, S7 and S3 indicate septin subunit names. **(F)** Laser scanning confocal imaging of hydrated SEPT2-GFP/6/7/3 filaments. Scale bar is 5 *μ*m.

Interpreting sedimentation coefficients requires an understanding of particle shape. By assembling existing and predicted structural models, we constructed a model of the SEPT2/6/7/3 hetero-octamer complex with coiled-coil regions (Figure 2B, C and Supplemental Figure 4). This was used to estimate the diffusion coefficient for the complex.^69^ Using these methods, we also modeled a 1:1:1:1 tetramer of SEPT2/SEPT6/SEPT7/SEPT3 (Figure 2D, E) and a SEPT2/SEPT6/SEPT7 hexamer (Supplemental Figure 4C). We note that these models combine multiple septin structural models^37,40,70–72^ by aligning structures using the hetero-trimer structure as a template^37^ and includes models for the coiled-coil regions that are computationally derived.^71,73^ The computationally derived parallel coiled-coil model is consistent with FRET between the SEPT6 and SEPT7 C-termini.^43^ In addition, the length of the coiled-coil model (~16 nm) is consistent with the 15-25 nm ‘paired filament’ separation observed for yeast septins.^62,74^ The resulting model should reasonably capture the coarse features that impact hydrodynamics.

From the convex hull around the model, we estimated the diffusion coefficient for the object including a hydration shell.^69^ By combining this diffusion coefficient with the molecular weight predicted for the complex, we estimated the sedimentation coefficient via the Svedberg equation. We predict a sedimentation coefficient of 9.7 S at 20°C in water for the SEPT2/6/7/3 hetero-octamer, and 6.9 S for the tetramer. Notably, the coiled-coils could plausibly adopt a more compact shape resulting in less hydrodynamic drag and faster sedimentation. To approximate an upper “compact limit” for the complex, we estimated the sedimentation coefficient for the complex without coiled-coils to be 13.0 S and 8.8 S for the SEPT2/6/7/3 hetero-octamer and tetramer, respectively. Realistically, the complex could not be this compact, instead this reflects an upper limit to the possible sedimentation coefficient range. As the 10.6 S average for the SEPT2/6/7/3 complex is well within the 9.7 −13.0 S range predicted for the octamer, we conclude that the complex is an octamer, albeit one that is somewhat more compact than the model shown in Figure 2B and C.

To test if these octamers can form filaments, we dialyzed the SEPT2-GFP/6/7/3 complex (Supplemental Figure 2B) into a reduced salt buffer (50 mM potassium chloride, 20 mM HEPES/NaOH pH 7.0, 2 mM MgCl_2_, 2 mM DTT) to induce filament formation.^17,75^ These samples were loaded into flow chambers compatible with imaging and observed by laser scanning confocal microscopy while still hydrated. Under these conditions, filament clusters are clearly observed (Figure 2F). Thus, our septin hetero-octamer complexes are functional with respect to filament formation.

### Hetero-octamers do not assemble from purified SEPT3 and SEPT2/6/7 complexes

A long-standing proposed model of the hetero-octamer places the SEPT3 subgroup members on the ends of a rod-like assembly, with the SEPT2/SEPT6/SEPT7 (called SEPT2/6/7 hereafter) hetero-hexamer as the inner six subunits (Figure 3A).^41,46^ If this model is correct, the hetero-octamer might be assembled by adding SEPT3 to pre-existing hetero-hexamer (Figure 3A). Purified SEPT2/6/7 hexamer was mixed with purified MBP-SEPT3 and then sedimented into a sucrose gradient. The MBP-SEPT3 fusion was left intact to allow ready discernment of SEPT2 and SEPT3 and does not interfere with complex formation (Figure 1C, D and Supplemental Figure 2B). MBP-SEPT3 clearly sediments with the other septins in the coexpressed complex (Figure 3B). When purified MBP-SEPT3 is mixed with purified hexamer, there was no change in sedimentation compared to isolated MBP-SEPT3 (Figure 3C vs 3E). Moreover, the SEPT2/6/7 sedimentation was not altered by addition of MBP-SEPT3 (Figure 3D vs. 3E). This indicates that once complexes are assembled, they do not easily combine with other septins. One plausible mechanism for this comes from yeast septin complexes, where GTP hydrolysis is thought to play a role in locking-down subunit exchange.^76^ Importantly, we conclude that producing complexes with all four septin sub-family members is most effective through co-expression.

**Figure 3:**
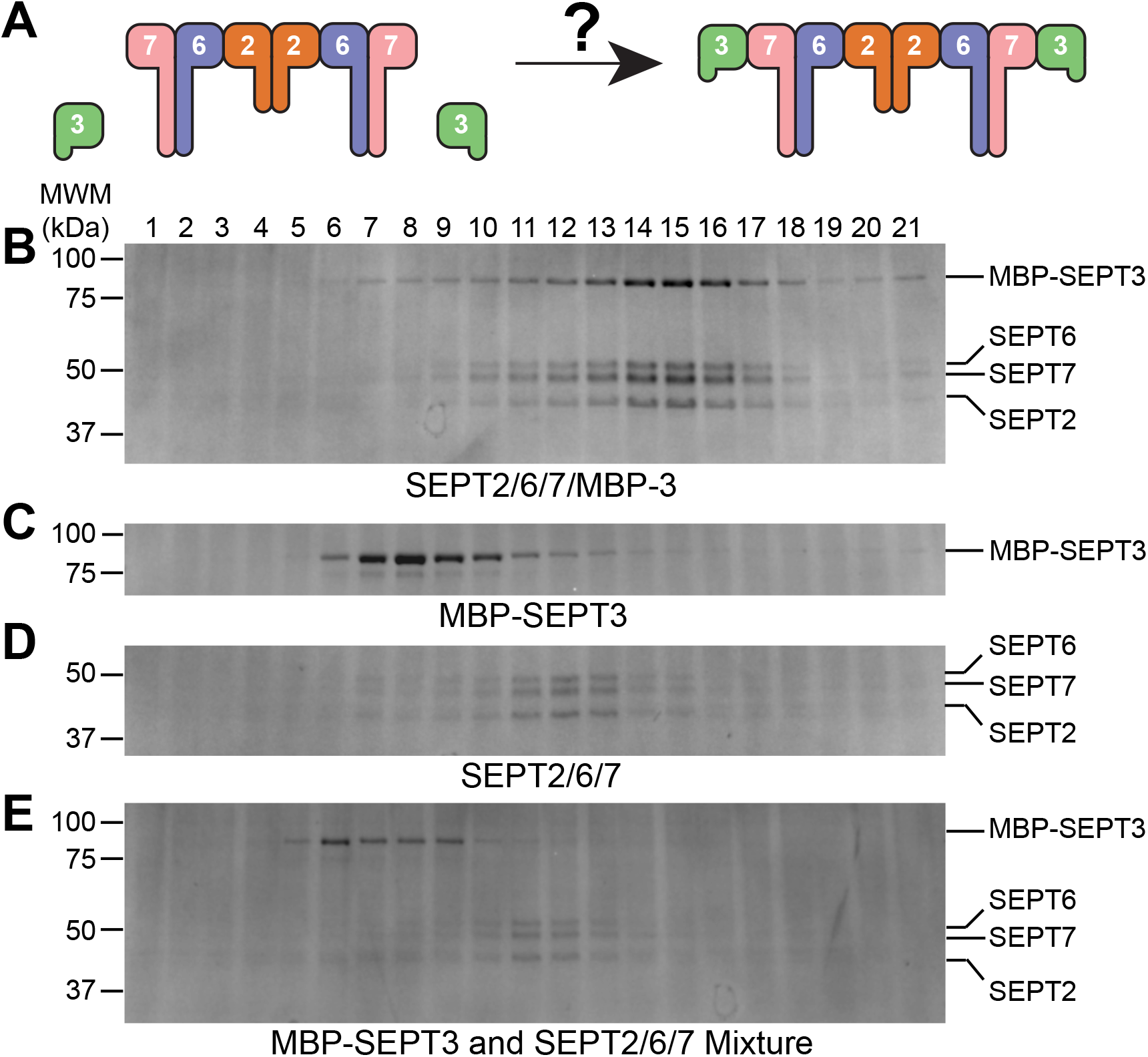
Post-purification assembly of SEPT2/6/7/3 octamer fails. **(A)** A long-standing model of septin octamer organization suggests that post-purification octamer assembly should be possible. **(B-E)** SDS-PAGE analysis of sucrose gradients of septin mixtures (CBB stained). Fractions are numbered from the top of the gradient (top fraction is #1). Mobilities of relevant species are indicated at the right and the mobilities of molecular weight markers at left. Gradients analyzing: **(B)** Co-expressed and purified SEPT2/SEPT6/SEPT7/MBP-SEPT3, **(C)** MBP-SEPT3, **(D)** SEPT2/SEPT6/SEPT7 and **(E)** post-purification mixture of SEPT2/6/7 with MBP-SEPT3

### Octamer assembly and stability are consistent with SEPT3 at an internal position

The inability to assemble the complex from purified materials is also consistent with a recent insight into the organization of the octamer. While the SEPT3 subunits had long been thought to occupy the terminal position of the octamer complex (Figure 4A, left) and SEPT7 to occupy the terminal positions of the hexamer complex,^37,46^ recent studies have called this into question.^45,47^ In the ‘new model’ of the octamer, SEPT2 is at the terminal positions of both complexes^45^ and SEPT3 occupies the central positions in the octamer (Figure 4A, right).^47^ The new octamer model is consistent with the failure to assemble complexes from purified SEPT2/6/7 hexamer and SEPT3 as SEPT3 would need to have the hexamer components disassembled and then re-assembled around it (Figure 3). While this observation can be explained multiple ways, it prompted us to explore the octamer organization further.

**Figure 4:**
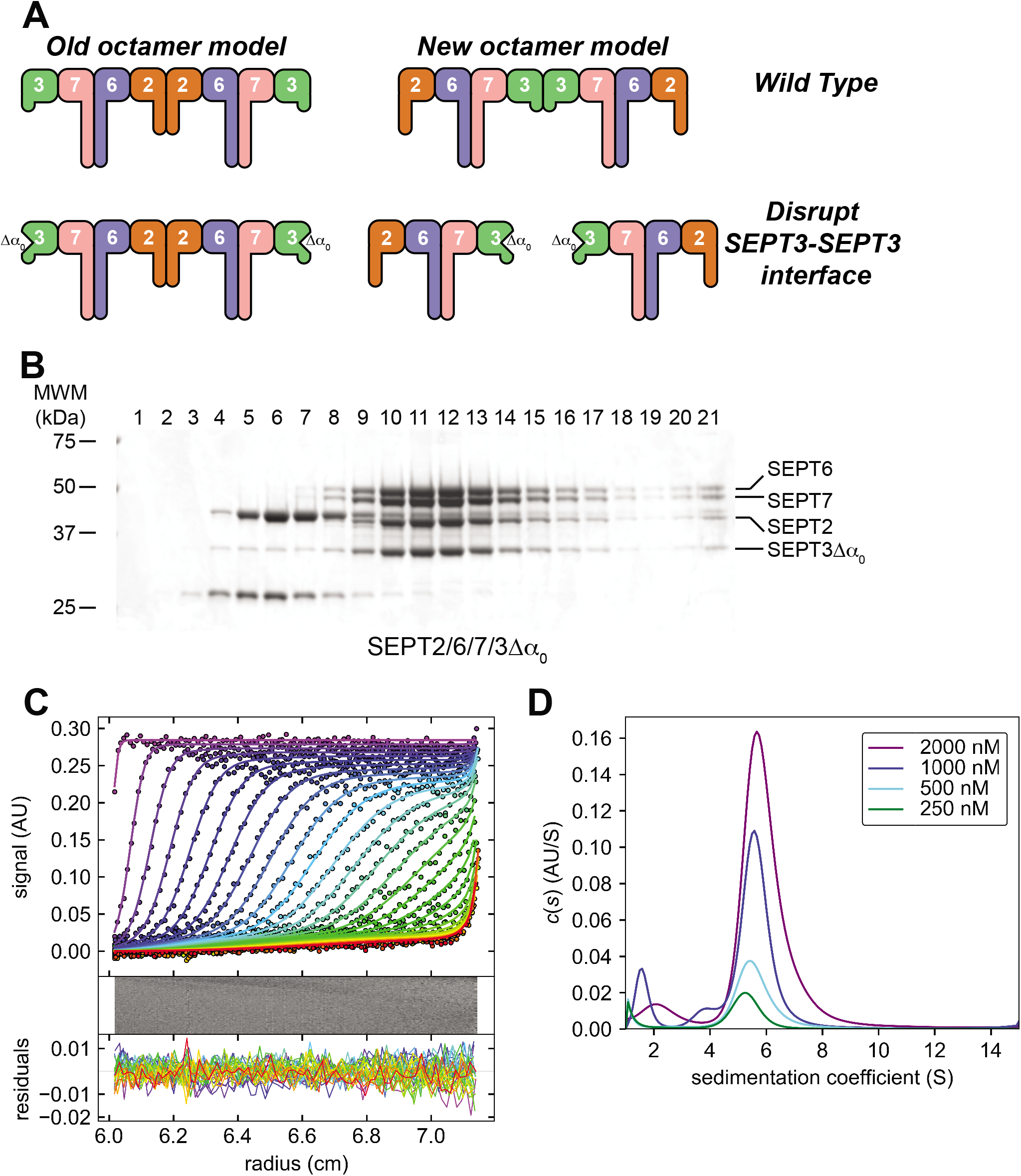
SEPT3Δα_0_ deletion disrupts the complex. **(A)** Schematic cartoons of proposed organizations for the octamer complex, with predicted impact of disrupting the SEPT3-SEPT3 interface through the SEPT3Δα_0_ truncation. **(B)** Sucrose gradient of SEPT2/6/7/3Δα_0_ with relevant mobilities at right and molecular weight markers at left. **(C)** Sedimentation velocity analytical ultracentrifugation (SV-AUC) of purified SEPT2/6/7/3Δα_0_ at 2000 nM. Circles are every third data point from every third scan. Colored lines are fit to data, with purple to blue for earlier time points and yellow to red for late time points. Middle and bottom subpanels are bitmap of residuals and residuals for each scan, respectively. **(D)** c(s) distributions for SEPT2/6/7/3Δα_0_ complex at the indicated concentrations.

If we assume this new octamer model is correct, disruption of the SEPT3-SEPT3 interface should break the octamer into tetramers (Figure 4A, right). Conversely, if the old octamer model is correct, SEPT3 is not in the middle of the complex and we should see little impact on complex sedimentation if we disrupt the SEPT3-SEPT3 interface (Figure 4A, left). To disrupt the interface, we deleted the N-terminus of SEPT3, including an alpha-helical structure element, α_0_, to produce SEPT3Δα_0_. By field convention, α_0_ is numbered to maintain consistency with the small GTPase fold septins evolved from. Based on expected similarities to other septins, this should disrupt the SEPT3-SEPT3 ‘N-/C-interface’ at the center of the new octamer model. This deletion prevents septin filament formation in several contexts.^37,38,40,46,77^ Moreover, while fulllength isolated SEPT3 adopts a heterogeneous mix of oligomeric states, this truncated SEPT3 was monomeric in the absence of GTP.^65^ Conveniently, the SEPT3Δα_0_ subunit resolves from SEPT2 in SDS-PAGE analysis, allowing identification of all four subunits in the resulting complex. The purified SEPT2/SEPT6/SEPT7/SEPT3Δα_0_ complex, called SEPT2/6/7/3Δα_0_ hereafter, still sediments as a unit with all four subunits readily identified (Figure 4B). Using our gradient calibration, this modified complex sedimented at 8.7 S. This falls outside of the expected range for octamers and near the compact extreme value predicted for the tetramer (8.8 S). Given this is an extreme value for tetramer sedimentation, we instead hypothesize that this slower sedimentation might reflect a dynamic equilibrium between tetramer and octamer species.

To further analyze the sedimentation of the SEPT2/6/7/3Δα_0_ complex we used SV-AUC, fitting the time course of sedimentation to a *c*(*s*) distribution (Figure 4C). This analysis found the SEPT3Δα_0_ complex sediments more slowly at lower concentrations, decreasing from a *s*_20,water_ value of 6.6 S (*s*_average_ = 6.0 S) at the highest concentration of complex to a value of 5.9 S (*s*_average_ = 5.3 S) at the lowest concentration. This concentration dependent change in sedimentation coefficient is a clear indication of dynamic changes in the complex stoichiometry. If we assume the lowest concentration sample approaches that of the smallest stable assembly, we can interpret the observed sedimentation coefficient. The *s*_20,water_ value of 5.9 S is actually a little below that predicted for our tetramer model (Figure 4D). This slower sedimentation could be explained in a few possible ways. First, in addition to complex mass, sedimentation is sensitive to compaction of the complex, where for a fixed protein mass, compact spheres sediment the fastest and less compact deviations from this sediment more slowly. This ‘slower than predicted sedimentation’ could be explained if the tetramer is a less compact structure, perhaps due to the SEPT2 coiled-coil region being disordered, in contrast to how it was modeled. For example, the SEPT2 coiled-coil could adopt extended conformations that point away from the plane defined by the GTPase domain rod and SEPT6/SEPT7 coiled-coils. Alternatively, the ‘slower than predicted sedimentation’ may be due to the tetramer disassembling into monomers, dimers and trimers that sediment more slowly due to reduced mass of the sub-complexes.

To exclude the alternative ‘disassembling tetramer’ possibility, we ran sucrose gradients at a range of concentrations and analyzed the fractions by SDS-PAGE (Supplemental Figure S5A-C). Using our gradient calibration, the sedimentation coefficient similarly decreased with concentration and approached a value of ~6 S in sucrose gradients, which is quite similar to the value of 5.9 S obtained using SV-AUC (Supplemental Figure S5D). Here, we observe that all members of the complex co-sediment (Supplemental Figure S5A-C), and thus reject the alternative hypothesis that the tetrameric complex is disassembling. Complex disassembly would not be expected to result in consistent sedimentation of all four subunits at near the predicted tetramer sedimentation rates. Thus, we feel the most likely explanation is that we are observing a tetramer-octamer equilibrium. The limiting lower sedimentation coefficient is consistent with a less compact tetramer structure and sedimentation coefficients between the tetramer and octamer values are due to dynamic formation and breakdown of the octamer. Thus, we conclude that the SEPT2/6/7/3Δα_0_ complex has a weakened SEPT3-SEPT3 interface affinity. This would place SEPT3 subunits at the center of the complex, consistent with the new complex model.^45,47^ Moreover, in this new model SEPT3 would need to associate with SEPT7 prior to formation of the hexamer, consistent with the observed lack of post-purification assembly (Figure 3).

This leaves open the question as to why the SEPT3Δα_0_ truncation merely weakens the SEPT3-SEPT3 N-/C- interface instead of fully disrupting it. Recent crystallographic observations identified the SEPT3 subgroup septins as having a modified N-/C- interface with two stable interaction modes, one of which excludes α_0_ from the interface.^40^ Thus, by deleting α_0_, we have disrupted only one of two possible interfaces, explaining why we have weakened as opposed to fully disrupted the interaction. Intriguingly, the preference for one interface over another appears to be GTP dependent.^40^. It is tempting to speculate that the SEPT3-SEPT3 interface may change in stability upon GTP hydrolysis, perhaps providing a mechanism for filament disassembly and complex recycling. Additionally, SEPT3 in isolation undergoes GTP dependent dimerization, although this has been attributed to dimerization via the G-interface instead of the N-/C- interface.^65,78^ This remains an interesting avenue for future research.

### Do septin octamer and hexamer complexes co-polymerize?

Finally, we asked if the octameric complexes co-assemble with hexamers into mixed filaments. In crystal structures, pairs of septin monomers interact via surfaces that include the GTP binding sites from both monomers (the ‘G-interface) or via surfaces that include the amino- and carboxy termini from both monomers (N-/C-interfaces), but there have been no observations of the G-interface of one septin interacting with the N-/C-interface of another. If the “old model” of the hexamer and octamer complexes is correct, then the hexamer and octamer complexes have the SEPT7 G- and SEPT3 N-/C- interfaces at their ends, respectively (Figure 5B, left). With the “new model”, both complexes have the N-/C- interface of SEPT2 at the ends (Figure 5B, right). Thus, the “old model” is inconsistent with hexamer and octamer coassembly, while the “new model” predicts co-assembly into mixed filaments via the SEPT2 N- /C- interface. As the slower sedimentation of our SEPT2/6/7/3Δα_0_ complex and the lack of octamer assembly from purified materials were consistent with the new assembly model, we looked for evidence of co-assembly between the complexes. We used the SEPT2-GFP tagged version of the SEPT2/6/7/3 octamer in conjunction with a SEPT2-mCherry tagged SEPT2/6/7 hexamer. Under our imaging conditions, each complex could individually produce clusters of filaments, with no bleed through between imaging channels (Figure 5B-G). When octamer and hexamer were co-polymerized, filament clusters were observed with clear intensity from both complexes (Figure 5H-J). From this we conclude that our recombinant octamer complex can polymerize into mixed filaments with recombinant hexamer.

**Figure 5:**
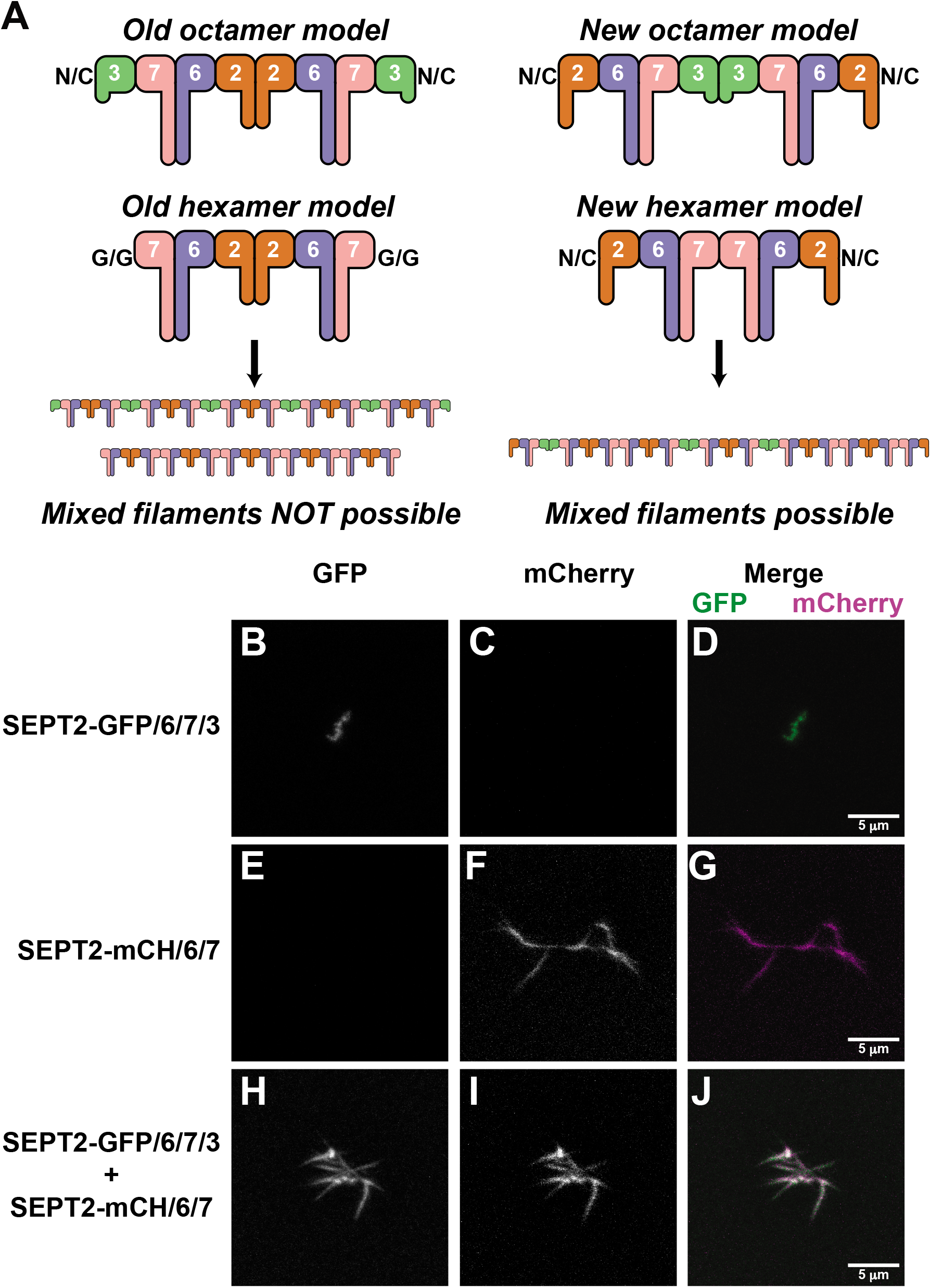
SEPT2/6/7/3 octamer and SEPT2/6/7 hexamer form mixed filaments. **(A)** Cartoon schematic how different octamer assembly models predict different results upon mixing of septin octamers and hexamers. **(B, C, and D)** SEPT2-GFP/6/7/3 octamer filaments formed as in Figure 2F, but imaged in GFP and mCherry channels, with merge, respectively. **(E, F, and G)** SEPT2-mCherry/6/7 hexamer filaments formed as in (Figure 2A), imaged in GFP and mCherry channels, with merge, respectively. **(H, I, and J)** SEPT2-GFP/6/7/3 and SEPT2-mCherry/6/7 complexes were mixed prior to dialysis to form filaments. GFP, mCherry and merge images shown, respectively. Scale bar is 5 *μ*m.

## Conclusions

Here we describe the first recombinant expression and purification strategy for mammalian septin complexes of defined composition that contain all four septin subgroups. These complexes are octamers in solution, consistent with that found for endogenous septins.^41,42^ In addition, this novel material is polymerization competent, reconstituting a primary biochemical activity of septins. Notably, the yield of complex is modest, with approximately 0.5 mg recovered per 6 L bacterial culture volume. This reflects a final volume of ~3 ml at ~1 μM octamer complex. This concentration is on the order of estimates of SEPT7 concentration in mammalian cells (see Supplemental Information in Ref. 79, suggesting that we can study biochemical behavior at cellular concentrations in future quantitative assays. We hope that this new complex provides an important tool for future biochemical study of septin assembly. Supporting this hope, this new protocol has already allowed us to confirm a new model of soluble septin complex assembly and to show that the complex forms mixed filaments with mammalian septin hexamers (Figure 5B). A similar observation of mixed octamer and hexamer filaments has been made with SEPT9 containing complexes.^47^ This new protocol opens the door to future studies of septin assembly including: how septin octamers interact with membranes and other cytoskeletal systems and how co-assembly with hexamers alters septin filament organization.

## Materials and Methods

### Vector construction

The expression plasmid for MBP-SEPT3/His6-SEPT2 was produced by inserting an expression cassette for human SEPT3 and mouse SEPT2, with an intervening Shine-Dalgarno/ribosome binding sequence and new start codon, into a modified pMAL2c parent vector (New England Biolabs, Ipswich, MA, USA). The modified vector had a TEV MBP cleavable fusion and a modified MCS. This vector was a gift from Michael K. Rosen. Using assembly cloning,^80^ septin sequences were inserted as a PCR fragment of SEPT3 excluding the 3 prime region, a PCR fragment of SEPT2 excluding the 5 prime region and a synthetic fragment consisting of the 3 prime of SEPT3, the intervening ribosome binding site and start codon and a TEV protease cleavable His6-tagged 5p region of SEPT2. A SEPT2-GFP fusion variant was produced by using PCR to remove the 3p stop codon from SEPT2 and to produce a GFP insert with an overlap to both the 3p end of SEPT2 and the pMAL2c derived vector, allowing fusion construct production through assembly cloning. The GFP insert was amplified from the plasmid cfSGFP2-N, which was a gift from Ikuo Wada (Addgene plasmid # 37533; http://n2t.net/addgene:37533; RRID:Addgene_37533).^81^ The co-expression vector for SEPT3Δα_0_/SEPT2 was produced by PCR amplifying the portion of SEPT3 used in assembly cloning but lacking the first 57 amino acids from SEPT3 (NCBI reference sequence: NP_663786), and then proceeding as in full-length co-expression construct construction. Plasmids pCDF-SEPT6/7 (co-expression plasmid encoding SEPT6 and SEPT7-StrepII),^63^ pET15b-SEPT2 (mouse SEPT2 with a Thrombin cleavable N-terminal His6 fusion)^63^ and pET28-SEPT3 (used for PCR amplification in construction of SEPT3/SEPT2 co-expression vector)^82^ were previously described. pET15b-SEPT2 was also modified to include a C-terminal fusion to mCherry, pET15b-SEPT2-mCh.

Blue fluorescent protein standards were produced by fusing sequence encoding BFP to MBP in the above modified version of pMAL2c or a further modified pMAL vector encoding a tandem fusion of MBP (referred to as 2MBP or diMBP). BFP inserts were PCR amplified from pNCS BFP, which was a gift from Erik Rodriguez & Roger Tsien^83^ (Addgene plasmid # 91757; http://n2t.net/addgene:91757; RRID:Addgene_91757). A C-terminal fusion of BFP to p97 in a pET15 derived expression vector was produced using standard PCR and restriction cloning methods; p97 vector was a gift from Patrick J. Loll^84^.

### Septin complex purification

Vectors for co-expressing MBP-SEPT3/His6-SEPT2 and SEPT6/SEPT7-StrepII were cotransformed into BL21(DE3)-T1^R^ cells and plated onto LB-Strep/Amp agar plates. Cultures grown at 37°C to an OD_600_ of ~0.8 were induced with 1 mM IPTG and allowed to express for three hours at 37°C. Cultures were harvested by centrifugation and resuspended in Lysis Buffer (500 mM Sodium Chloride, 50 mM Tris pH 8.0, 5 mM MgCl_2_, 2 mM EGTA, 2 mM DTT, 1mM PMSF). Cell suspensions were snap frozen in liquid nitrogen and stored at −80°C until needed. Cells were lysed by extrusion (Emulsiflex C5, Avestin Inc. Ottawa Canada), and centrifuged at 17,000 rpm in a JA-20 rotor at 4°C for 40 minutes. Clarified lysates were bound to amylose beads and washed extensively with AmyWB (500 mM Sodium Chloride, 20 mM Tris pH 8.0, 5 mM MgCl_2_), and eluted with AmyEB (500 mM Sodium Chloride, 20 mM Tris pH 8.0, 5 mM MgCl_2_, 30 mM Maltose). Amylose elution pool was bound to Nickel NTA (NiNTA) affinity beads (Qiagen, Germantown MD, USA), washed with NiWB (500 mM Sodium Chloride, 20 mM Tris pH 8.0, 5 mM MgCl_2_, 10 mM Imidazole) and eluted using NiEB (500 mM Sodium Chloride, 20 mM Tris pH 8.0, 5 mM MgCl_2_, 250 mM Imidazole). Elutions with the highest protein content were pooled and digested overnight with TEV protease. The digested sample was further purified using a sucrose density gradient.

MBP-SEPT3 was expressed using only the MBP-SEPT3/His6-SEPT2 vector. The purification method remained the same as for the octamer through the amylose step. The NiNTA step was the same as for the octamer purification, except the flow through was retained. The NiNTA flow through was clarified by centrifugation at 50,000 rpm in a Type 70 Ti rotor at 4°C for 1 hour. The supernatant was applied to a Superdex 200 pg 26/600 column (GE) and developed with a buffer of 500 mM Sodium Chloride, 20 mM Tris pH 8.0, 5 mM MgCl_2_, 2 mM EGTA, 1 mM DTT. Fractions were assessed by SDS-PAGE. Samples were aliquoted into <100 uL samples, snap frozen in liquid nitrogen and stored at −80°C.

The SEPT2/6/7 complex was co-expressed as for the octamer but using the pET15b-SEPT2 and pCDF-SEPT6/SEPT7-StrepII vectors.^63^ The complex was purified by NiNTA (same buffers as for the octameric). Filaments were formed by dialysis into 100 mM potassium chloride, 10 mM HEPES/NaOH pH 7.0, 1 mM MgCl_2_, 0.5 mM EGTA, 1 mM DTT. Filaments were collected using centrifugation in a Type 70 Ti rotor, spinning at 55,000 rpm for two hours, at 4°C. Filaments were dissolved into complexes by resuspending in a high salt buffer (500 mM Sodium Chloride, 50 mM Tris pH 8.0, 5 mM MgCl_2_, 1 mM DTT), then centrifuged at 16,000*g for 20 minutes at 4°C. The supernatant was aliquoted, snap frozen in liquid nitrogen and stored at −80°C. SEPT2-mCh/6/7 complex was purified by similar methods but expressed from the pET15b-SEPT2-mCh and pCDF-SEPT6/SEPT7-StrepII vectors.

### Sucrose gradients

Preparative and analytic sucrose gradients used the same method. 10% and 30% weight to volume sucrose buffers (otherwise containing 500 mM Potassium Chloride, 50 mM Tris pH 8.0, 5 mM MgCl_2_, 2 mM EGTA, 2 mM DTT) were prepared and filtered. Gradients were prepared by layering 2.25 mL of 10% sucrose buffer on top of 2.25 mL of 30% sucrose buffer and rotating at an angle to produce a linear gradient, as assessed by tracer dye concentration. 500 *μL* samples were loaded onto the gradient immediately centrifuged at 50,000 rpm for 14 hours in a SW-55 rotor at 4°C. Samples were collected by hand into 250 microliter fractions. Fractions were analyzed using SDS-PAGE and bands detected using ‘Stain Free’ or Coomassie brilliant blue. Gel imaging was accomplished with GelDoc EZ with white or ‘Stain Free’ imaging plates (BioRad, Hercules CA USA). Following preparative gradient purification, samples were pooled, aliquoted into <100 uL samples, snap frozen in liquid nitrogen, and stored at −80°C. For analytic sucrose gradients, samples were dialyzed into sucrose free buffer prior to loading.

### Modeling of hydrodynamic parameters

Models SEPT2/6/7/3 tetramers and octamers used PDB ID #2QAG (SEPT2/SEPT6/SEPT7 trimer^37^ as the starting point. The trimer has a SEPT2-SEPT6 G-/G- interface and a SEPT6/SEPT7 N-/C- interface. To place the SEPT3 subunit, we used the ‘super’ command in PyMol (Schrödinger, LLC) to align a copy of 2QAG SEPT2 onto SEPT7. The copy of SEPT6 post alignment was used to place SEPT3 (4Z51).^40^ G-domains for SEPT2 and SEPT7 were updated by aligning PDB# 2QNR^72^ and 6N0B^70^, respectively. α_0_ of the SEPT2 subunit from 2QAG was kept associated with SEPT2 (PDB ID: 2QNR).^72^ Thus, a SEPT2/6/7/3 tetramer G-domain backbone was produced. To model the coiled-coils, which are absent from the G-domain structures and from 2QAG, we built a model for the C-terminal SEPT6-SEPT7 CC hetero-dimer and SEPT2 homo-dimer using CCfold.^71^ These were built slightly outside of the identified coiled-coil region^73^ such that we could extend the final a-helix in the SEPT7 G-domain to orient the coiled-coil model. To complete the octamer, two tetramers were assembled end-to-end via their SEPT3 surfaces using the lattice contacts between 6N0B chains A and D as a template.^70^ Hexamers were similarly constructed and models without coiled-coils were produced by deleting the atoms for those regions from the above models. The generated PDB files were then analyzed using HullRad^69^ to find hydrodynamic radius and diffusion coefficients. The diffusion coefficient along with the predicted complex molecular weight and the Svedberg equation was used to estimate the sedimentation coefficients for the different complexes. The convex hull display was accomplished in PyMol using a script provided by the authors of HullRad.

### Analytical Ultracentrifugation

SEPT2/6/7/3Δα_0_ complex samples were loaded into 1.2 cm thick, two-channel Epon centerpieces, fitted with quartz windows (Beckman USA). Reference sectors were filled with the buffer that samples were dialyzed against (500 mM Potassium chloride, 50 mM Tris/HCl pH 8.0, 5 mM MgCl_2_, 2 mM EGTA, 200 μM TCEP and 20 mM HEPES/NaOH pH 7.0). Concentration was estimated by UV absorbance (e_290nm_ = 70,678 and 69,709 M^−1^cm^−1^ for SEPT2/6/7/3 wild type and SEPT2/6/7/3Δα_0_ complexes, respectively) and diluted to the indicated concentrations using the dialysis buffer (concentrations are in 1:1:1:1 tetramer units). Sample cells were loaded into an An-50 Ti or An-60 Ti rotor and spun at 40,000 rpm at 20°C, monitoring sedimentation by A_280_. BFP tagged constructs at 2 to 5.5 μM were dialyzed against 100 mM Sodium Chloride with 5 mM HEPES pH 7.0. SV-AUC experiments were run on the BFP materials following absorbance at 380 nm (near the BFP absorbance peak) and 280 nm, in separate experiments. SV-AUC data was analyzed using SEDFIT^85^ to construct *c*(*s*) distributions, with optimization of the frictional ratio, meniscus and cell bottom. Buffer parameters were estimated using SEDNTERP.^86^ *s*_20,water_ was determined in SEDFIT by integrating the *c*(*s*) peak and correcting for buffer effects. Data-fit-residual plots and c(s) overlays were produced using GUSSI.^87^

### Filament imaging

Septin samples were prepared under high salt conditions (500 mM potassium chloride, 20 mM HEPES/NaOH pH 7.0, 2 mM MgCl_2_, 2 mM DTT). Samples were: 1.2 μM SEPT2-GFP/6/7/3 (100% GFP labeled), 8.5 μM SEPT2/6/7-STREPII with 0.5 μM SEPT2-mCherry/6/7, and a one-to-one mix of the GFP and mCherry labeled samples. Samples were loaded into 50 microliter micro-dialysis buttons, overlaid with dialysis membrane (SpectraPOR 14 kDa MWCO) and dialyzed for ~16 hours at 4°C against a 400-fold excess of imaging buffer (50 mM potassium chloride, 20 mM HEPES/NaOH pH 7.0, 2 mM MgCl_2_, 2 mM DTT). 18 and 25 mm square glass coverslips (#1.5, VWR Scientific) were cleaned using multiple sonication steps in 1% Micro-90 detergent, 100% ethanol and 1 M potassium hydroxide, washing with ultra-pure water following each step. Coverslips were dried under a stream of argon and assembled into flow chambers using ~80 μm thick double stick tape. BSA (Gold Biotechnology) was labeled with NHS-LC-Biotin (Pierce Biotechnology). Flow chambers were initially wet with ultrapure water and then 1 mg/ml biotinylated BSA was flowed into to the chamber using surface tension and incubated for 10 minutes at room temperature. The chamber was washed twice with imaging buffer. 0.1 mg/ml streptavidin in the imaging buffer was flowed in and bound for 10 minutes before washing twice with imaging buffer. Finally, dialyzed septin filament samples were flowed into the chamber and the chamber sealed with nail polish. Septin filaments were immobilized through collective action of many weak-binding events of the SEPT7-Strep II tag to streptavidin on the chamber surface. Samples were kept in the dark and imaged the same day they were prepared. Samples were imaged using an FV3000 laser scanning confocal (Olympus), with 1.35 n.a. 100x UPLSAPO objective with silicone oil immersion media. GFP was excited with a 488 nm laser and detected by emission at 500-540 nm. mCherry was excited at 561 nm and detected by emission at 570-620 nm. To reduce bleed through, excitation lasers (488 nm and 561 nm) were excited in separate phases. Ten vertical sections were acquired over 3.9 μm. Images were processed using FIJI/ImageJ^88–90^ using a max Z projection followed by contrast adjustment such that intensities were matched between images to be compared.

---

**Supplementary Figure 1:**
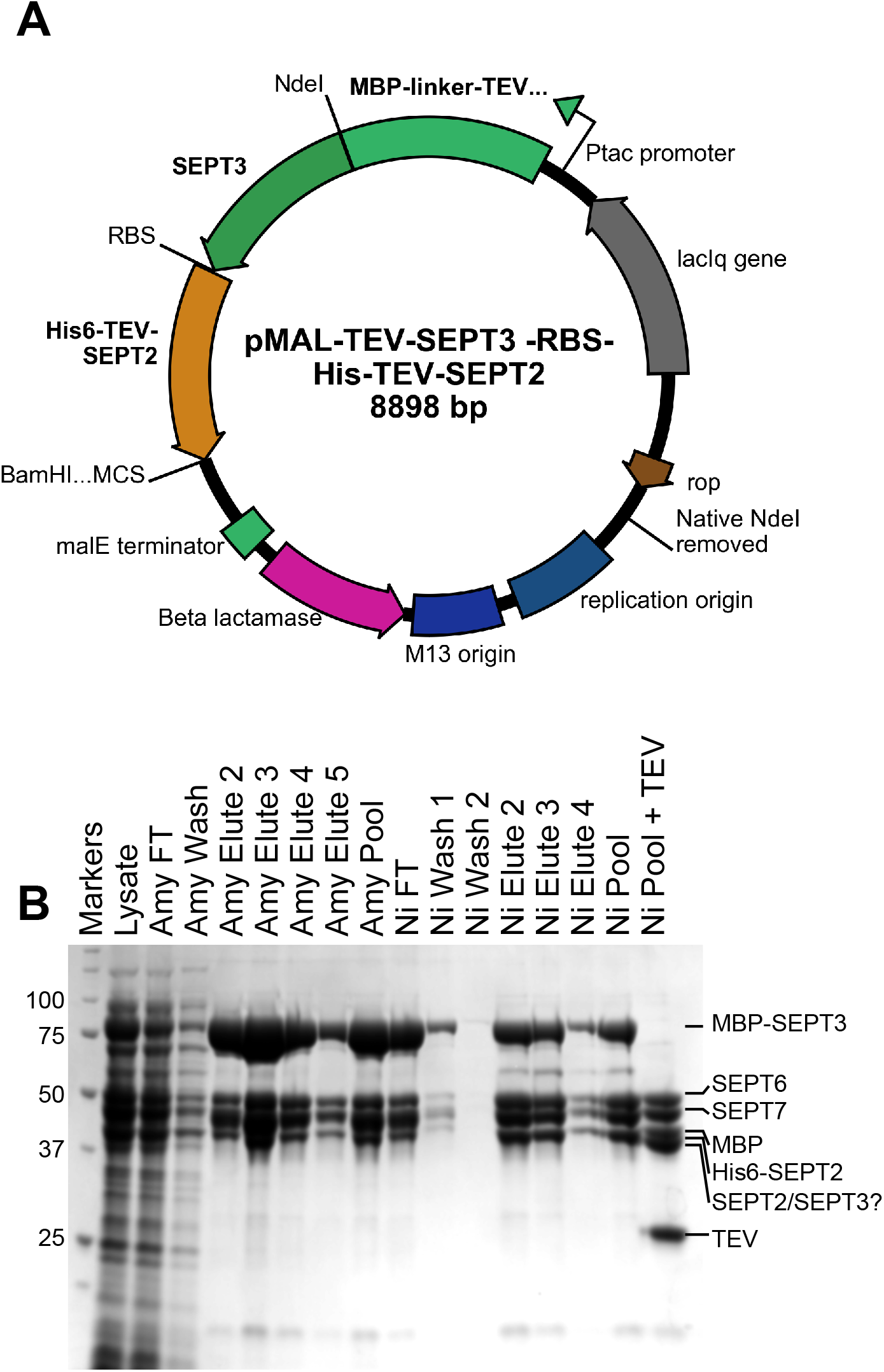
Production of SEPT2/6/7/3 complex. **(A)** Plasmid map of expression vector for MBP-SEPT3 and His6-SEPT2. **(B)** SEPT2/6/7/3 affinity purification gel analysis, CBB stained. Amylose (Amy) and Ni-NTA agarose (Ni) affinity steps purify for MBP fusions and His6 fusions, respectively. Flow through (FT) from a step and elution fractions plus elution pool are shown. Molecular weight marker mobilities are shown at left and relevant product mobilities shown at right. “Ni Pool + TEV” shows proteins purified after TEV protease digestion, note loss of MBP-SEPT3 band. A similar change in mobility for His6-SEPT2 is obscured by the MBP released from MBP-SEPT3.

**Supplementary Figure 2:**
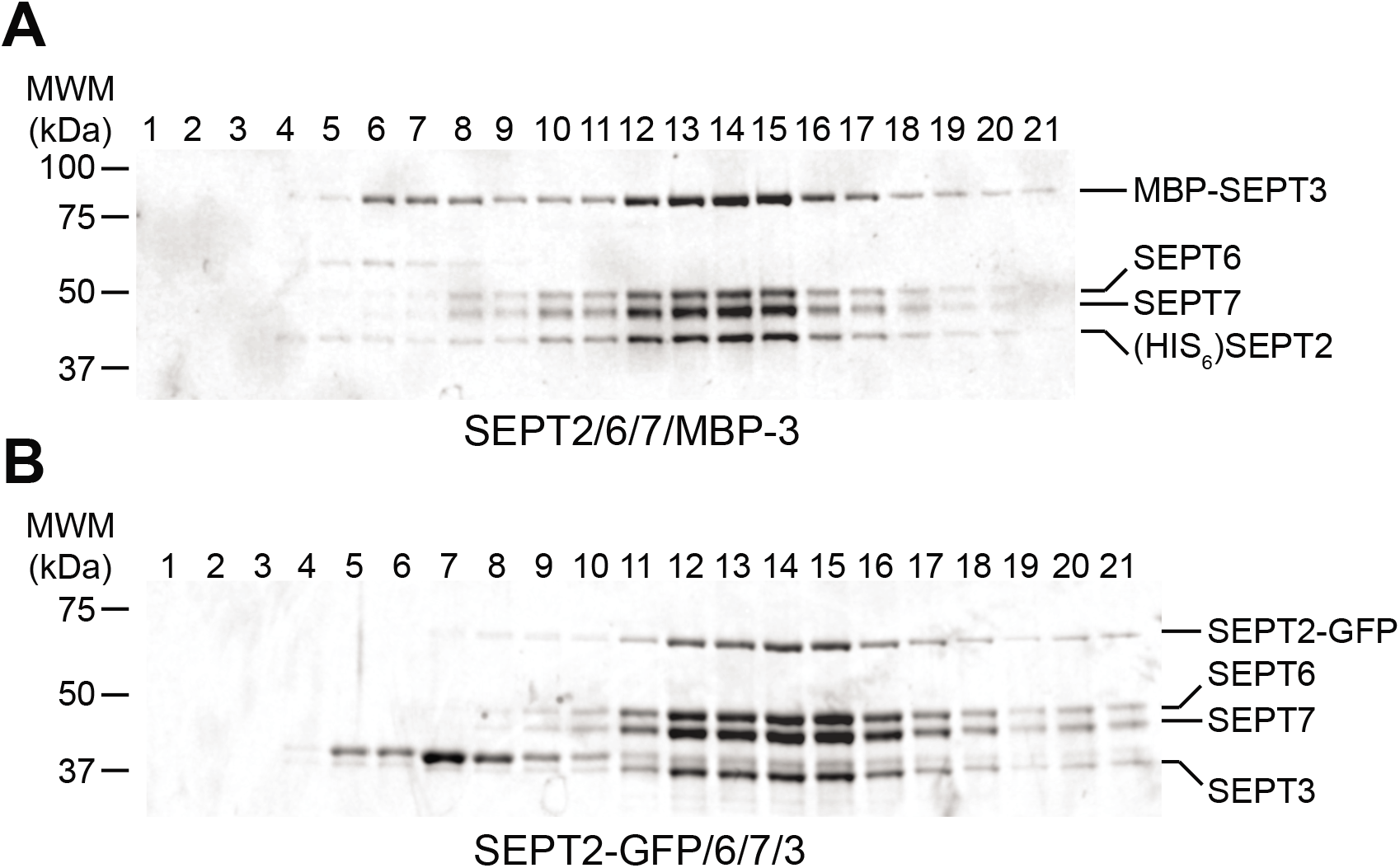
SEPT2-GFP/6/7/3 and SEPT2/6/7/MBP-3 sediment as stable complexes. Sucrose gradient sedimentation of septin complexes with tags: **(A)** SEPT2-GFP/6/7/3 complex and **(B)** SEPT2/6/7/MBP-3 complex (same as in Figure 1, but without TEV digestion). Relevant mobilities shown at right and molecular weight markers at left.

**Supplementary Figure 3:**
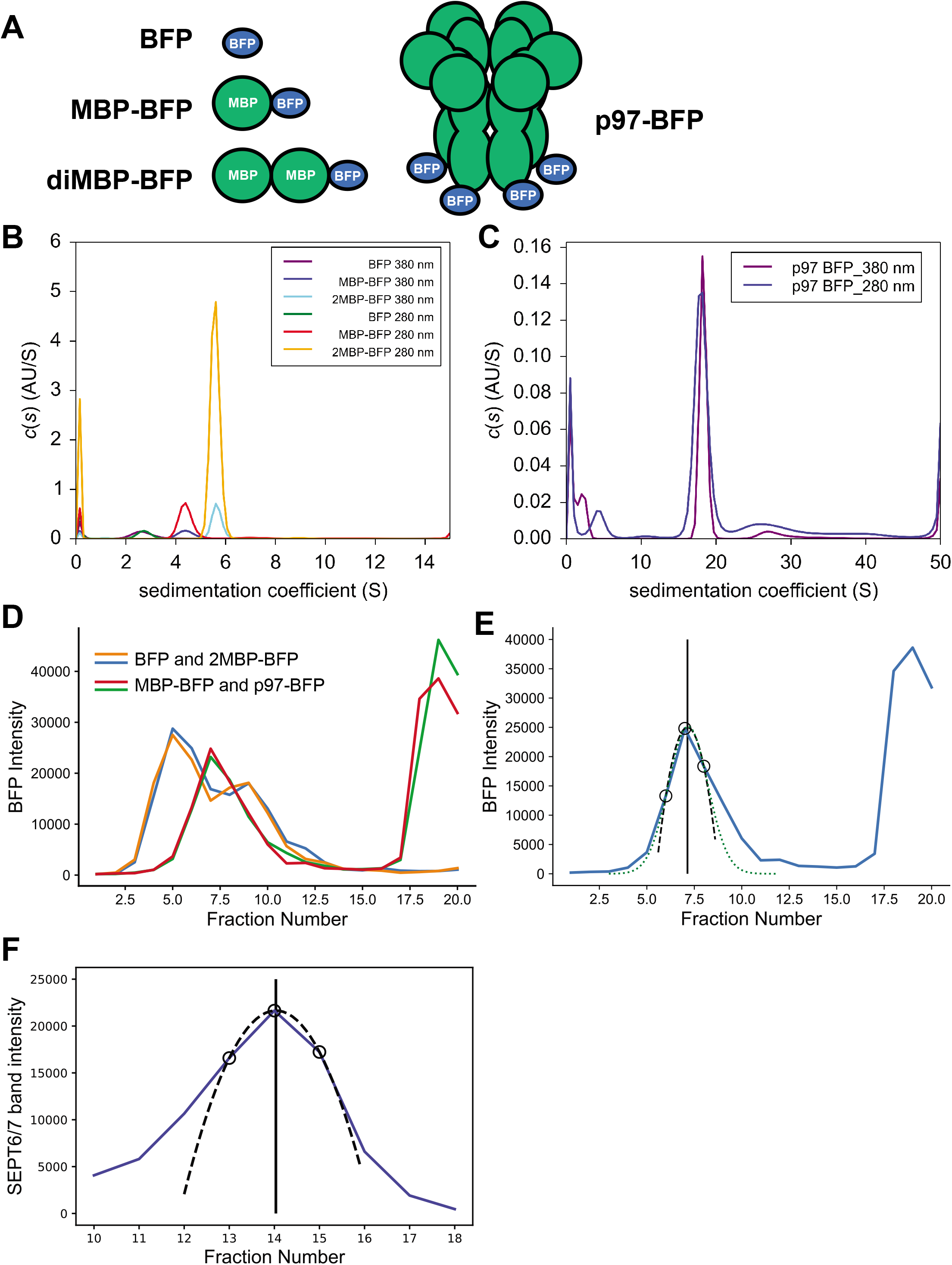
Calibration of sucrose gradients. **(A)** A series blue fluorescent protein (BFP) fusions were produced. These were purified and analyzed by SV-AUC. **(B and C)** c(s) distributions from analysis of BFP standards detecting at 280 nm or 380 nm to detect aromatic amino acids or BFP, respectively. Principle peaks were integrated and corrected for buffer effects to find *s*_20°C, water_. **(D)** BFP fluorescence from fractions of sucrose gradients separating mixtures of BFP and 2MBP-BFP or MBP-BFP and p97-BFP, with each mixture in duplicate. **(E)** To better quantify the sedimentation, the peak fraction and one from either side were used to fit a parabola and the center was used as the ‘peak fraction’ value. Choice of a parabola was not essential, fitting to a Guassian curve yielded an estimate of the center within 1% of the parabola estimate. The parabolic center estimates were used in Figure 2A. **(F)** To quantify the sedimentation for septin complexes, ‘Stain Free’ bands were integrated, and a similar procedure to that in (E) was used. (E and F) Blue curve: connecting intensities for each fraction. Open circles: data points used for parabola estimate. Dashed line: fit parabola. Vertical solid line: center of parabola. Green dots: fit Guassian curve.

**Supplementary Figure 4:**
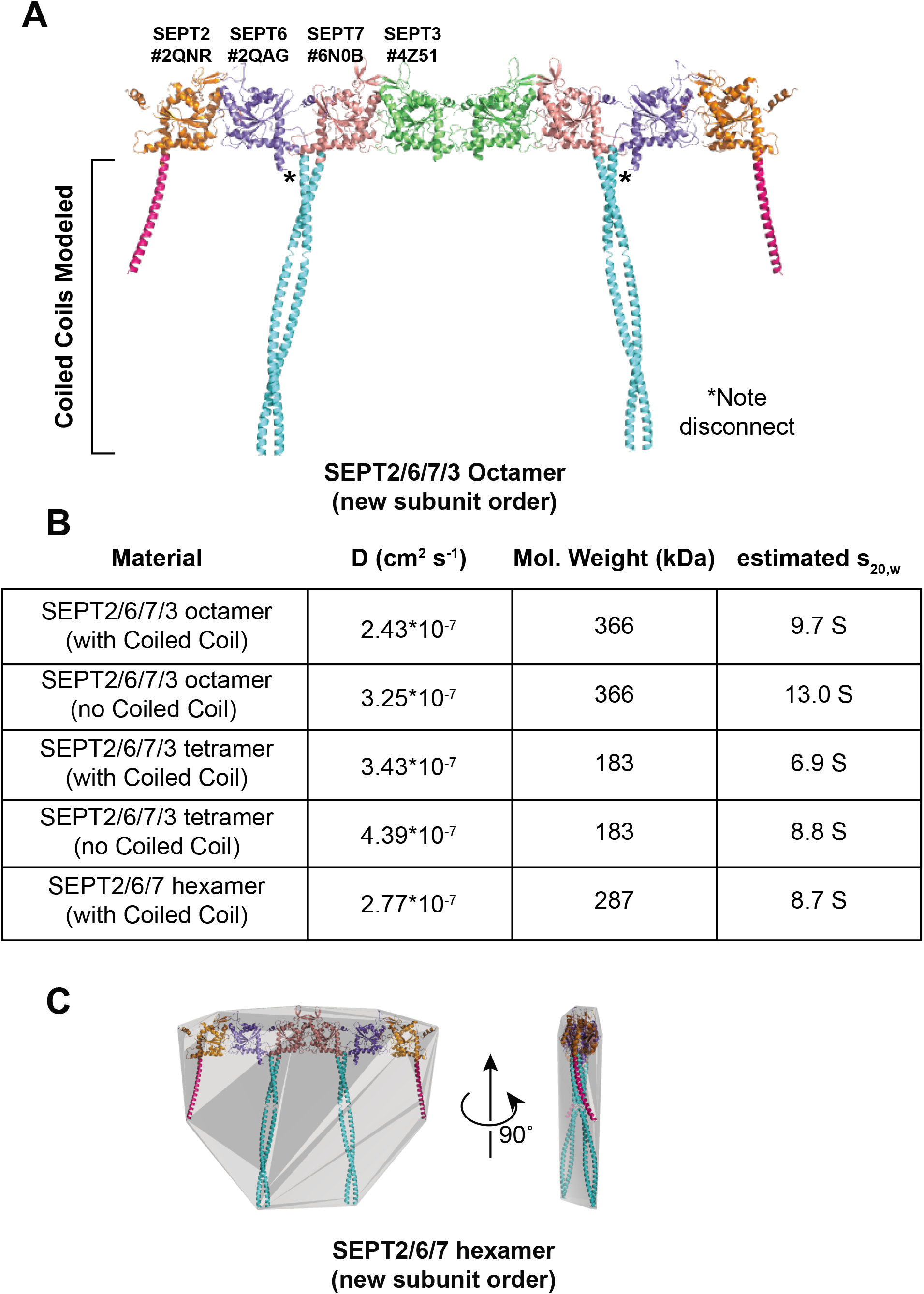
Estimation of sedimentation for septin complex models. **(A)** SEPT2/6/7/3 octamer model with sub-model origins indicated. **(B)** Table of models and diffusion coefficients, D, estimated using HullRad. In addition, the molecular weight estimated for the complex (using the sequence predicted from the expression construct) and the estimated sedimentation coefficient under standard conditions are provided. **(C)** Model for the SEPT2/6/7 hexamer with modeled coiled-coils (cyan and magenta) and convex hull for estimation of diffusion coefficients.

**Supplementary Figure 5:**
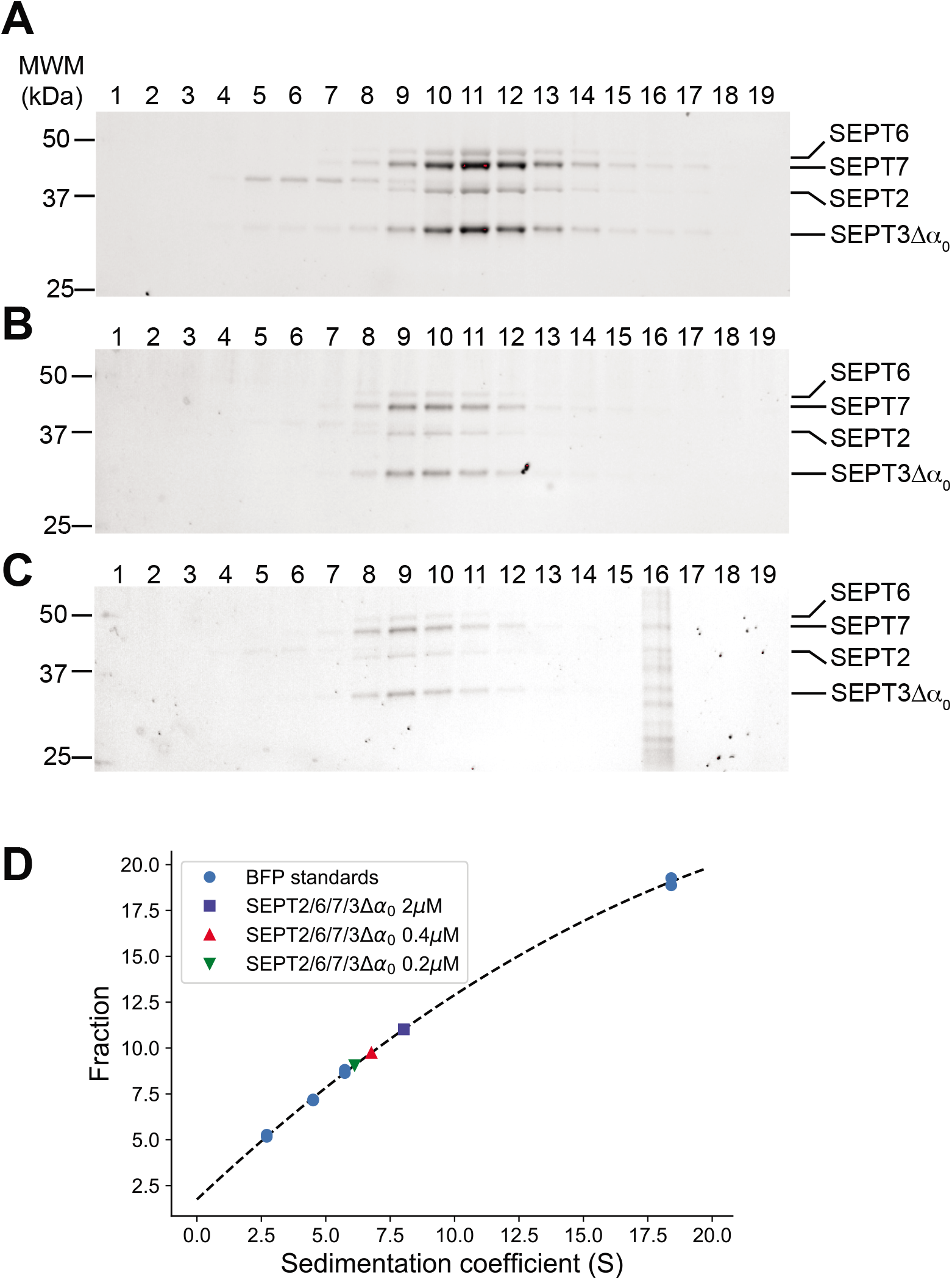
Sedimentation of SEPT2/6/7/3Δα_0_ complex at different concentrations. The sedimentation of the SEPT2/6/7/3Δα_0_ complex decreases with decreasing concentration. Sedimentation was measured using sucrose gradients and analyzed using SDS-PAGE imaged using ‘Stain Free’ methods. Diluted SEPT2/6/7/3Δα_0_ materials were loaded at 2000 nM **(A)**, 400 nM **(B)** and 200 nM **(C)**. Notably, all four subunits co-sediment for each sample, but the sedimentation slows at lower concentrations (peak fractions are at lower fraction numbers). (C) Lane 16 has an unexplained contaminant that is not expected to originate from the gradient itself (gradients do not generally produce peaks one fraction wide). The imaging exposure time used in panels (A)-(C) is not the same due to the 10-fold range in concentrations loaded. **(D)** Sucrose gradient calibrations were used to identify the sedimentation coefficient for each sample above (see symbols in the in-panel legend).

## Acknowledgements

Shae Padrick is supported by startup funds from the Drexel University College of Medicine and the Department of Biochemistry and Molecular Biology. E.T.S. is supported by NIH/NIGMS grants 5RO1 GM097664-9 and 1R35 GM136337. We thank Robert Fairman and Ronen Marmorstein for access to analytical ultracentrifugation instrumentation. We thank Dr. Patrick Loll for critical reading of the manuscript and the p97 vector. We thank Meagan Tomasso for a critical reading of the manuscript and the SEPT2/6/7 complex. We thank the College of Medicine and the Department of Pharmacology and Physiology access to the Olympus FV3000 laser scanning confocal microscope.

## References

1 Hartwell, L. H., Culotti, J., Pringle, J. R. & Reid, B. J. Genetic control of the cell division cycle in yeast. Science 183, 46–51, doi:10.1126/science.183.4120.46 (1974).

2 Haarer, B. K. & Pringle, J. R. Immunofluorescence localization of the Saccharomyces cerevisiae CDC12 gene product to the vicinity of the 10-nm filaments in the mother-bud neck. Mol Cell Biol 7, 3678–3687, doi:10.1128/mcb.7.10.3678 (1987).

3 Pan, F., Malmberg, R. L. & Momany, M. Analysis of septins across kingdoms reveals orthology and new motifs. BMC Evol Biol 7, 103, doi:10.1186/1471-2148-7-103 (2007).

4 Estey, M. P., Di Ciano-Oliveira, C., Froese, C. D., Bejide, M. T. & Trimble, W. S. Distinct roles of septins in cytokinesis: SEPT9 mediates midbody abscission. J Cell Biol 191, 741–749, doi:10.1083/jcb.201006031 (2010).

5 Karasmanis, E. P. et al. A Septin Double Ring Controls the Spatiotemporal Organization of the ESCRT Machinery in Cytokinetic Abscission. Curr Biol 29, 2174–2182 e2177, doi:10.1016/j.cub.2019.05.050 (2019).

6 Ribet, D. et al. SUMOylation of human septins is critical for septin filament bundling and cytokinesis. J Cell Biol 216, 4041–4052, doi:10.1083/jcb.201703096 (2017).

7 Mostowy, S. & Cossart, P. Septins: the fourth component of the cytoskeleton. Nat Rev Mol Cell Biol 13, 183–194, doi:10.1038/nrm3284 (2012).

8 Saarikangas, J. & Barral, Y. The emerging functions of septins in metazoans. EMBO Rep 12, 1118–1126, doi:10.1038/embor.2011.193 (2011).

9 Spiliotis, E. T., Hunt, S. J., Hu, Q., Kinoshita, M. & Nelson, W. J. Epithelial polarity requires septin coupling of vesicle transport to polyglutamylated microtubules. J Cell Biol 180, 295–303, doi:10.1083/jcb.200710039 (2008).

10 Beites, C. L., Xie, H., Bowser, R. & Trimble, W. S. The septin CDCrel-1 binds syntaxin and inhibits exocytosis. Nat Neurosci 2, 434–439, doi:10.1038/8100 (1999).

11 Beites, C. L., Campbell, K. A. & Trimble, W. S. The septin Sept5/CDCrel-1 competes with alpha-SNAP for binding to the SNARE complex. Biochem J 385, 347–353, doi:10.1042/BJ20041090 (2005).

12 Huang, Y. W. et al. Mammalian septins are required for phagosome formation. Mol Biol Cell 19, 1717–1726, doi:10.1091/mbc.E07-07-0641 (2008).

13 Caudron, F. & Barral, Y. Septins and the lateral compartmentalization of eukaryotic membranes. Dev Cell 16, 493–506, doi:10.1016/j.devcel.2009.04.003 (2009).

14 Hu, Q. et al. A septin diffusion barrier at the base of the primary cilium maintains ciliary membrane protein distribution. Science 329, 436–439, doi:10.1126/science.1191054 (2010).

15 Yamada, S. et al. Septin Interferes with the Temperature-Dependent Domain Formation and Disappearance of Lipid Bilayer Membranes. Langmuir 32, 12823–12832, doi:10.1021/acs.langmuir.6b03452 (2016).

16 Dolat, L. et al. Septins promote stress fiber-mediated maturation of focal adhesions and renal epithelial motility. J Cell Biol 207, 225–235, doi:10.1083/jcb.201405050 (2014).

17 Kinoshita, M., Field, C. M., Coughlin, M. L., Straight, A. F. & Mitchison, T. J. Self- and actin-templated assembly of Mammalian septins. Dev Cell 3, 791–802 (2002).

18 Joo, E., Surka, M. C. & Trimble, W. S. Mammalian SEPT2 is required for scaffolding nonmuscle myosin II and its kinases. Dev Cell 13, 677–690, doi:10.1016/j.devcel.2007.09.001 (2007).

19 Calvo, F. et al. Cdc42EP3/BORG2 and Septin Network Enables Mechano-transduction and the Emergence of Cancer-Associated Fibroblasts. Cell Rep 13, 2699–2714, doi:10.1016/j.celrep.2015.11.052 (2015).

20 Kremer, B. E., Adang, L. A. & Macara, I. G. Septins regulate actin organization and cellcycle arrest through nuclear accumulation of NCK mediated by SOCS7. Cell 130, 837–850, doi:10.1016/j.cell.2007.06.053 (2007).

21 Kim, J. & Cooper, J. A. Septins regulate junctional integrity of endothelial monolayers. Mol Biol Cell 29, 1693–1703, doi:10.1091/mbc.E18-02-0136 (2018).

22 Torraca, V. & Mostowy, S. Septins and Bacterial Infection. Front Cell Dev Biol 4, 127, doi:10.3389/fcell.2016.00127 (2016).

23 Mostowy, S. et al. Entrapment of intracytosolic bacteria by septin cage-like structures. Cell Host Microbe 8, 433–444, doi:10.1016/j.chom.2010.10.009 (2010).

24 Krokowski, S. et al. Septins Recognize and Entrap Dividing Bacterial Cells for Delivery to Lysosomes. Cell Host Microbe 24, 866–874 e864, doi:10.1016/j.chom.2018.11.005 (2018).

25 Pfanzelter, J., Mostowy, S. & Way, M. Septins suppress the release of vaccinia virus from infected cells. J Cell Biol 217, 2911–2929, doi:10.1083/jcb.201708091 (2018).

26 Mostowy, S. et al. Septins regulate bacterial entry into host cells. PLoS One 4, e4196, doi:10.1371/journal.pone.0004196 (2009).

27 Scholz, R. et al. Novel Host Proteins and Signaling Pathways in Enteropathogenic E. coli Pathogenesis Identified by Global Phosphoproteome Analysis. Mol Cell Proteomics 14, 1927–1945, doi:10.1074/mcp.M114.046847 (2015).

28 Bai, X. et al. Novel septin 9 repeat motifs altered in neuralgic amyotrophy bind and bundle microtubules. J Cell Biol 203, 895–905, doi:10.1083/jcb.201308068 (2013).

29 Peterson, E. A. & Petty, E. M. Conquering the complex world of human septins: implications for health and disease. Clin Genet 77, 511–524, doi:10.1111/j.1399-0004.2010.01392.x (2010).

30 Connolly, D. et al. Septin 9 amplification and isoform-specific expression in peritumoral and tumor breast tissue. Biol Chem 395, 157–167, doi:10.1515/hsz-2013-0247 (2014).

31 Angelis, D. & Spiliotis, E. T. Septin Mutations in Human Cancers. Front Cell Dev Biol 4, 122, doi:10.3389/fcell.2016.00122 (2016).

32 Pous, C., Klipfel, L. & Baillet, A. Cancer-Related Functions and Subcellular Localizations of Septins. Front Cell Dev Biol 4, 126, doi:10.3389/fcell.2016.00126 (2016).

33 Warren, J. D. et al. Septin 9 methylated DNA is a sensitive and specific blood test for colorectal cancer. BMC Med 9, 133, doi:10.1186/1741-7015-9-133 (2011).

34 Kinoshita, M. The septins. Genome Biol 4, 236, doi:10.1186/gb-2003-4-11-236 (2003).

35 Kinoshita, M. Assembly of mammalian septins. J Biochem 134, 491–496, doi:10.1093/jb/mvg182 (2003).

36 Versele, M. & Thorner, J. Some assembly required: yeast septins provide the instruction manual. Trends Cell Biol 15, 414–424, doi:10.1016/j.tcb.2005.06.007 (2005).

37 Sirajuddin, M. et al. Structural insight into filament formation by mammalian septins. Nature 449, 311–315, doi:10.1038/nature06052 (2007).

38 Sirajuddin, M., Farkasovsky, M., Zent, E. & Wittinghofer, A. GTP-induced conformational changes in septins and implications for function. Proc Natl Acad Sci U S A 106, 16592–16597, doi:10.1073/pnas.0902858106 (2009).

39 Valadares, N. F., d’ Muniz Pereira, H., Ulian Araujo, A. P. & Garratt, R. C. Septin structure and filament assembly. Biophys Rev 9, 481–500, doi:10.1007/s12551-017-0320-4 (2017).

40 Castro, D. K. S. d. V. et al. A complete compendium of crystal structures for the human SEPT3 subgroup reveals functional plasticity at a specific septin interface. IUCrJ 7, 462–479, doi:10.1107/S2052252520002973 (2020).

41 Sellin, M. E., Sandblad, L., Stenmark, S. & Gullberg, M. Deciphering the rules governing assembly order of mammalian septin complexes. Mol Biol Cell 22, 3152–3164, doi:10.1091/mbc.E11-03-0253 (2011).

42 Sellin, M. E., Stenmark, S. & Gullberg, M. Cell type-specific expression of SEPT3-homology subgroup members controls the subunit number of heteromeric septin complexes. Mol Biol Cell 25, 1594–1607, doi:10.1091/mbc.E13-09-0553 (2014).

43 Low, C. & Macara, I. G. Structural analysis of septin 2, 6, and 7 complexes. J Biol Chem 281, 30697–30706, doi:10.1074/jbc.M605179200 (2006).

44 Neubauer, K. & Zieger, B. The Mammalian Septin Interactome. Front Cell Dev Biol 5, 3, doi:10.3389/fcell.2017.00003 (2017).

45 Mendonca, D. C. et al. A revised order of subunits in mammalian septin complexes. Cytoskeleton (Hoboken) 76, 457–466, doi:10.1002/cm.21569 (2019).

46 Kim, M. S., Froese, C. D., Estey, M. P. & Trimble, W. S. SEPT9 occupies the terminal positions in septin octamers and mediates polymerization-dependent functions in abscission. J Cell Biol 195, 815–826, doi:10.1083/jcb.201106131 (2011).

47 Soroor, F. et al. Revised subunit order of mammalian septin complexes explains their in vitro polymerization properties. bioRxiv, doi:10.1101/569871 (2019).

48 Ihara, M. et al. Cortical organization by the septin cytoskeleton is essential for structural and mechanical integrity of mammalian spermatozoa. Dev Cell 8, 343–352, doi:10.1016/j.devcel.2004.12.005 (2005).

49 Smith, C. et al. Septin 9 Exhibits Polymorphic Binding to F-Actin and Inhibits Myosin and Cofilin Activity. J Mol Biol 427, 3273–3284, doi:10.1016/j.jmb.2015.07.026 (2015).

50 Spiliotis, E. T. Spatial effects – site-specific regulation of actin and microtubule organization by septin GTPases. J Cell Sci 131, doi:10.1242/jcs.207555 (2018).

51 Omrane, M. et al. Septin 9 has Two Polybasic Domains Critical to Septin Filament Assembly and Golgi Integrity. iScience 13, 138–153, doi:10.1016/j.isci.2019.02.015 (2019).

52 Tanaka-Takiguchi, Y., Kinoshita, M. & Takiguchi, K. Septin-mediated uniform bracing of phospholipid membranes. Curr Biol 19, 140–145, doi:10.1016/j.cub.2008.12.030 (2009).

53 Nakos, K., Rosenberg, M. & Spiliotis, E. T. Regulation of microtubule plus end dynamics by septin 9. Cytoskeleton (Hoboken) 76, 83–91, doi:10.1002/cm.21488 (2019).

54 Karasmanis, E. P. et al. Polarity of Neuronal Membrane Traffic Requires Sorting of Kinesin Motor Cargo during Entry into Dendrites by a Microtubule-Associated Septin. Dev Cell 46, 204–218 e207, doi:10.1016/j.devcel.2018.06.013 (2018).

55 Nagata, K. et al. Filament formation of MSF-A, a mammalian septin, in human mammary epithelial cells depends on interactions with microtubules. J Biol Chem 278, 18538–18543, doi:10.1074/jbc.M205246200 (2003).

56 Garcia, G., 3rd et al. Assembly, molecular organization, and membrane-binding properties of development-specific septins. J Cell Biol 212, 515–529, doi:10.1083/jcb.201511029 (2016).

57 Garcia, G., 3rd et al. Subunit-dependent modulation of septin assembly: budding yeast septin Shs1 promotes ring and gauze formation. J Cell Biol 195, 993–1004, doi:10.1083/jcb.201107123 (2011).

58 Bridges, A. A. et al. Septin assemblies form by diffusion-driven annealing on membranes. Proc Natl Acad Sci U S A 111, 2146–2151, doi:10.1073/pnas.1314138111 (2014).

59 Cannon, K. S., Woods, B. L., Crutchley, J. M. & Gladfelter, A. S. An amphipathic helix enables septins to sense micrometer-scale membrane curvature. J Cell Biol 218, 1128–1137, doi:10.1083/jcb.201807211 (2019).

60 Booth, E. A., Vane, E. W., Dovala, D. & Thorner, J. A Forster Resonance Energy Transfer (FRET)-based System Provides Insight into the Ordered Assembly of Yeast Septin Hetero-octamers. J Biol Chem 290, 28388–28401, doi:10.1074/jbc.M115.683128 (2015).

61 Booth, E. A. & Thorner, J. A FRET-based method for monitoring septin polymerization and binding of septin-associated proteins. Methods Cell Biol 136, 35–56, doi:10.1016/bs.mcb.2016.03.024 (2016).

62 Bertin, A. et al. Saccharomyces cerevisiae septins: supramolecular organization of heterooligomers and the mechanism of filament assembly. Proc Natl Acad Sci U S A 105, 8274–8279, doi:10.1073/pnas.0803330105 (2008).

63 Mavrakis, M. et al. Septins promote F-actin ring formation by crosslinking actin filaments into curved bundles. Nat Cell Biol 16, 322–334, doi:10.1038/ncb2921 (2014).

64 Xue, J. et al. Septin 3 (G-septin) is a developmentally regulated phosphoprotein enriched in presynaptic nerve terminals. J Neurochem 91, 579–590, doi:10.1111/j.1471-4159.2004.02755.x (2004).

65 Macedo, J. N. et al. The structure and properties of septin 3: a possible missing link in septin filament formation. Biochem J 450, 95–105, doi:10.1042/BJ20120851 (2013).

66 Kim, M. S., Froese, C. D., Xie, H. & Trimble, W. S. Uncovering principles that control septin-septin interactions. J Biol Chem 287, 30406–30413, doi:10.1074/jbc.M112.387464 (2012).

67 Ladner, C. L., Yang, J., Turner, R. J. & Edwards, R. A. Visible fluorescent detection of proteins in polyacrylamide gels without staining. Analytical Biochemistry 326, 13–20, doi:10.1016/j.ab.2003.10.047 (2004).

68 Edwards, R. A., Jickling, G. & Turner, R. J. The light-induced reactions of tryptophan with halocompounds. Photochem Photobiol 75, 362–368, doi:10.1562/0031-8655(2002)075<0362:tlirot>2.0.co;2 (2002).

69 Fleming, P. J. & Fleming, K. G. HullRad: Fast Calculations of Folded and Disordered Protein and Nucleic Acid Hydrodynamic Properties. Biophys J 114, 856–869, doi:10.1016/j.bpj.2018.01.002 (2018).

70 Brognara, G., Pereira, H. M., Brandao-Neto, J., Araujo, A. P. U. & Garratt, R. C. Revisiting SEPT7 and the slippage of beta-strands in the septin family. J Struct Biol 207, 67–73, doi:10.1016/j.jsb.2019.04.015 (2019).

71 Guzenko, D. & Strelkov, S. V. CCFold: rapid and accurate prediction of coiled-coil structures and application to modelling intermediate filaments. Bioinformatics 34, 215–222, doi:10.1093/bioinformatics/btx551 (2018).

72 Human septin 2 in complex with GDP. To be published, doi:10.2210/pdb2qnr/pdb.

73 Ludwiczak, J., Winski, A., Szczepaniak, K., Alva, V. & Dunin-Horkawicz, S. DeepCoil-a fast and accurate prediction of coiled-coil domains in protein sequences. Bioinformatics 35, 2790–2795, doi:10.1093/bioinformatics/bty1062 (2019).

74 DeMay, B. S. et al. Septin filaments exhibit a dynamic, paired organization that is conserved from yeast to mammals. J Cell Biol 193, 1065–1081, doi:10.1083/jcb.201012143 (2011).

75 Frazier, J. A. et al. Polymerization of purified yeast septins: evidence that organized filament arrays may not be required for septin function. J Cell Biol 143, 737–749, doi:10.1083/jcb.143.3.737 (1998).

76 Weems, A. & McMurray, M. The step-wise pathway of septin hetero-octamer assembly in budding yeast. Elife 6, doi:10.7554/eLife.23689 (2017).

77 McMurray, M. A. et al. Septin filament formation is essential in budding yeast. Dev Cell 20, 540–549, doi:10.1016/j.devcel.2011.02.004 (2011).

78 Ortore, M. G. et al. Structural and Thermodynamic Properties of Septin 3 Investigated by Small-Angle X-Ray Scattering. Biophys J 108, 2896–2902, doi:10.1016/j.bpj.2015.05.015 (2015).

79 Hein, M. Y. et al. A human interactome in three quantitative dimensions organized by stoichiometries and abundances. Cell 163, 712–723, doi:10.1016/j.cell.2015.09.053 (2015).

80 Gibson, D. G. et al. Enzymatic assembly of DNA molecules up to several hundred kilobases. Nat Methods 6, 343–345, doi:10.1038/nmeth.1318 (2009).

81 Suzuki, T. et al. Development of cysteine-free fluorescent proteins for the oxidative environment. PLoS One 7, e37551, doi:10.1371/journal.pone.0037551 (2012).

82 Angelis, D., Karasmanis, E. P., Bai, X. & Spiliotis, E. T. In silico docking of forchlorfenuron (FCF) to septins suggests that FCF interferes with GTP binding. PLoS One 9, e96390, doi:10.1371/journal.pone.0096390 (2014).

83 Cubitt, A. B. et al. Understanding, improving and using green fluorescent proteins. Trends Biochem Sci 20, 448–455, doi:10.1016/s0968-0004(00)89099-4 (1995).

84 Rao, M. V., Williams, D. R., Cocklin, S. & Loll, P. J. Interaction between the AAA(+) ATPase p97 and its cofactor ataxin3 in health and disease: Nucleotide-induced conformational changes regulate cofactor binding. J Biol Chem 292, 18392–18407, doi:10.1074/jbc.M117.806281 (2017).

85 Schuck, P. Size-distribution analysis of macromolecules by sedimentation velocity ultracentrifugation and lamm equation modeling. Biophys J 78, 1606–1619, doi:10.1016/S0006-3495(00)76713-0 (2000).

86 Laue, T., Shah, B. V., Ridgeway, T. M. & Pelletier, S. L. Computer-aided interpretation of analytical sedimentation data for proteins. Chemistry, 90–125 (1992).

87 Brautigam, C. A. Calculations and Publication-Quality Illustrations for Analytical Ultracentrifugation Data. Methods Enzymol 562, 109–133, doi:10.1016/bs.mie.2015.05.001 (2015).

88 Schindelin, J. et al. Fiji: an open-source platform for biological-image analysis. Nat Methods 9, 676–682, doi:10.1038/nmeth.2019 (2012).

89 Schneider, C. A., Rasband, W. S. & Eliceiri, K. W. NIH Image to ImageJ: 25 years of image analysis. Nat Methods 9, 671–675, doi:10.1038/nmeth.2089 (2012).

90 Rueden, C. T. et al. ImageJ2: ImageJ for the next generation of scientific image data. BMC Bioinformatics 18, 529, doi:10.1186/s12859-017-1934-z (2017).

